# Integrated molecular and pharmacological characterization of patient-derived xenografts from bladder and ureteral cancers identifies new potential therapies

**DOI:** 10.1101/2022.04.19.488770

**Authors:** Hervé Lang, Claire Béraud, Luc Cabel, Jacqueline Fontugne, Myriam Lassalle, Clémentine Krucker, Florent Dufour, Clarice S. Groeneveld, Victoria Dixon, Xiangyu Meng, Aurélie Kamoun, Elodie Chapeaublanc, Aurélien De Reynies, Xavier Gamé, Pascal Rischmann, Ivan Bieche, Julien Masliah-Planchon, Romane Beaurepere, Yves Allory, Véronique Lindner, Yolande Misseri, François Radvanyi, Philippe Lluel, Isabelle Bernard-Pierrot, Thierry Massfelder

**Author notes:** To whom correspondence should be addressed: Dr Isabelle Bernard-Pierrot, Institut Curie, 26 rue d’Ulm, 75005 Paris, Dr Philippe Lluel, Urosphere, Canal Biotech II, 3 Rue des Satellites, 31400 Toulouse. These authors contributed equally to the work (Hervé Lang, Claire Béraud, Luc Cabel). these authors co-supervised the study (Isabelle Bernard-Pierrot, Thierry Massfelder).

## Abstract

**Background:** Muscle-invasive bladder cancer (MIBC) and upper urinary tract urothelial carcinoma (UTUC) are molecularly heterogeneous. Despite chemotherapies, immunotherapies or anti-FGFR treatments, these tumors are still of poor outcome. Our objective was to develop a bank of patient-derived xenografts (PDXs) recapitulating molecular heterogeneity of MIBC and UTUC, to facilitate preclinical identification of therapies.

**Methods:** Fresh tumors were obtained from patients and subcutaneously engrafted into immune-compromised mice. Patient tumors and matched PDXs were compared regarding histopathology, transcriptomic (microarrays) and genomic profiles (targeted-NGS). Several PDXs were treated with chemotherapy (cisplatin/gemcitabine) or targeted therapies (FGFR and EGFR inhibitors).

**Results:** 31 PDXs were established from 1 non-MIBC, 25 MIBC, 5 upper urinary tract tumors, including 28 urothelial (UCC) and 3 squamous-cell carcinomas (SCC). Integrated genomic and transcriptomic profiling identified PDXs of 3 different consensus molecular subtypes (Basal/Squamous, Luminal papillary and Luminal unstable), and included *FGFR3*-mutated PDXs. High histological and genomic concordance was found between matched patient tumor/PDX. Discordance in molecular subtypes, such as a basal/squamous patient tumor giving rise to a luminal papillary PDX, was observed (n=5) at molecular and histological levels. Ten models were treated with cisplatin-based chemotherapy and we did not observe association between subtypes and response. Of the 3 basal/squamous models treated with anti-EGFR therapy, two models were sensitive and one model, of sarcomatoid variant, was resistant. Treatment of 3 FGFR3-mutant PDXs with combined FGFR/EGFR inhibitors was more efficient than anti-FGFR3 treatment alone.

**Conclusions:** We developed preclinical PDX models that recapitulate the molecular heterogeneity of MIBCs and UTUC, including actionable mutations, which will represent an essential tool in therapy development. Pharmacological characterization of the PDXs suggested that upper urinary tract and MIBCs, UCC but also SCC, with similar molecular characteristics could benefit from the same treatments including anti-FGFR for FGFR3-mutated tumors and anti-EGFR for basal ones and showed a benefit for combined FGFR/EGFR inhibition in FGFR3-mutant PDXs, compared to FGFR inhibition alone.

## Introduction

Bladder cancer (BCa) is the 9th most common cancer type worldwide with an estimated 549,000 new cases in 2018 (1). Histologically, 90 to 95% of BCa are urothelial cell carcinomas (UCC) and 5% are squamous cell carcinomas (SCC) (2). Although less frequent, UCC may also develop in the upper urinary tract (2-5% of UCC). Muscle-invasive BCa (MIBC) are of very poor outcome, with an overall 5-year survival of 50-60% and less than 10% for patients with localized disease or distant metastasis, respectively. Localized UCC/SCC-MIBC and high-risk upper tract urothelial carcinoma (UTUC) are treated by radical cystectomy and radical nephroureterectomy respectively, with the addition of adjuvant or neoadjuvant chemotherapies (2,3). In advanced setting UCC, immune checkpoint or FGFR inhibitors (for tumors presenting *FGFR2/3* genetic alterations) may also be proposed (2,4), while there is no standard of care treatment for SCC. However, despite these treatments, the outcome remains poor, and identification of new therapies is still needed. The development of relevant pre-clinical models is critical to reach this objective.

MIBCs constitute a heterogeneous group of tumors at the morphological and molecular levels. With the aim to improve prediction of clinical outcome and treatment response, an international consortium has recently reached a consensus molecular classification based on transcriptomic data, for MIBC, including 6 subtypes, facilitating inter-study comparisons (5). These subtypes can be divided into broad luminal (differentiated) and basal/squamous (Ba/Sq) groups. Although current systemic treatments are not based on molecular classification, some subtype features are associated with treatment response. For example, Ba/Sq tumors have worse prognosis than luminal-papillary (LumP) tumors [5], which commonly show genetic alterations of FGFR3 (6). These subtypes have also been suggested to have a different sensitivity to chemotherapies, although results are currently inconsistent between studies (7–10). The Ba/Sq subtype and bladder SCC are associated with EGFR activation and sensitivity to anti-EGFR treatments in preclinical models (5,11–14). UTUC are molecularly comparable to MIBCs but have some distinct features, such as higher prevalence of microsatellite instability (MSI) (15).

The molecular heterogeneity of MIBCs and UTUC and their divergent therapy sensitivities entails the need for a representative panel of preclinical models for unbiased therapeutic evaluation. Therapeutic testing in patient-derived tumor xenografts (PDX) is highly effective in predicting the efficacy of both chemotherapies and targeted therapies (16,17). Although few studies have reported the development of PDX models from MIBC, none have performed an integrated genomic, transcriptomic and pharmacological characterization of a bank of PDX reflecting the biological diversity of MIBC (18–20). Recently, a bank of 17 PDXs from UTUC has been reported (21) but no model exists for SCC, for which no standard of care is established.

In the present report, we describe the development and characterization of a bank of 31 PDXs, which maintain the characteristics of patient tumors and reflect the diversity of the molecular subtypes of bladder and upper urinary tract cancers. Evaluation of pharmacological responses to standard of care and targeted therapies suggests that non-sarcomatoid basal/squamous tumors, notably SCC, could benefit from anti-EGFR therapies. In *FGFR3*-mutated tumor PDXs, an anti-FGFR and anti-EGFR combination therapy improves tumor response compared to FGFR inhibition alone.

## MATERIALS AND METHODS

### Animals

Four- to five-week-old immunodeficient mice (male) were purchased from Charles River Laboratories (L’Arbresle, France). Mice were maintained under specific pathogen-free conditions. Their care and housing were conducted in accordance with the European Community Council Directive 2010/63/UE and the French Ministry for Agriculture, Agrifood and Forestry Decree 2013-118. Experimental protocols were reviewed by CEEA-122 Ethical Committee for Protection of Animals used for Scientific Purposes and approved by French Ministry for National Education, Higher Education and Research under the number *APAFIS#14811-2018042316405732 v4*. The animal facility was maintained under artificial lighting (12 h) between 7:00 am to 7:00 pm in a controlled ambient temperature of 22 ± 2°C, and relative humidity rate maintained at 55 ± 10%.

### Specimen acquisition

From January 2009 to October 2019, patient tumors and matched normal tissues were obtained from 153 BCa patients undergoing surgery either at the Hospital of Strasbourg (France) (n=135) or the Hospital of Toulouse (France) (n=18) in accordance with all relevant guidelines and regulations. All patients provided written informed consent. Incoming material of every donor patient was anonymized by receiving a chronological unique number, subsequently used to identify the corresponding PDX model. The specimens were examined, sectioned, and selected by pathologists for histological analyses and xenografts. Clinical and demographic information were obtained prospectively.

### Patient-derived xenografts (PDX) establishment

PDX models of MIBC were generated by engrafting tumor tissues directly obtained from patients. Viable tumor tissue was macro-dissected and tumor pieces were then prepared for implantation. NMRI nude (Rj:NMRI-Foxn1^nu/nu^) immunodeficient mice strain was used for tissue implantation. Grafts were implanted into the interscapular fat pad. When s.c. xenograft tumors reached ∼ 1000-1500 mm3, they were serially transplanted for expansion into new mice. In addition, harvested xenograft material was cryopreserved for future implantations and/or fixed in 4% formalin for 24h before paraffin embedding and/or stored at -80°C for subsequent analyses. A model was defined as established when stable growth over at least three passages and regrowth after a freeze-thaw cycle could be observed. Take rate (proportion of mice developing tumors after transplantation of the PDX) and passage time were recorded for every model and every individual passage. Tumor growth was determined weekly by a two-dimensional measurement with a caliper. Tumor volume was calculated as: TV (mm3) = [length (mm) x width (mm)2]*π/6, where the length and width are the longest and shortest diameters of the tumor, respectively. Animals were sacrificed when tumor volume reached 2000 mm^3^.

### Histopathology and Immunohistochemistry

For all PDX models, primary and passaged tumors preserved in formalin for 24h were paraffin-embedded, sectioned into 4 μm-thick cuts and placed on glass slides. Analysis of hematoxylin and eosin (H&E) stained slides were performed by two experienced uropathologists.

### RNA/ DNA/Protein extraction

Each frozen PDX fragment was ground to powder and subdivided for triple RNA, DNA and protein extraction. RNA isolation was performed using trizol while DNA isolation used Phenol/Chloroform/Isoamyl Alcohol extraction. See “Western blot” section for protein extraction method.

### Real-time reverse transcription-quantitative PCR

Reverse transcription was performed on 1□µg of total RNA using a high-capacity cDNA reverse transcription kit (Thermo Fisher Scientific, Illkirch, France). cDNAs were amplified by PCR in a Roche real-time thermal cycler, with the Roche Taqman master mix (Roche) and Taqman probe/primer pairs as follows:

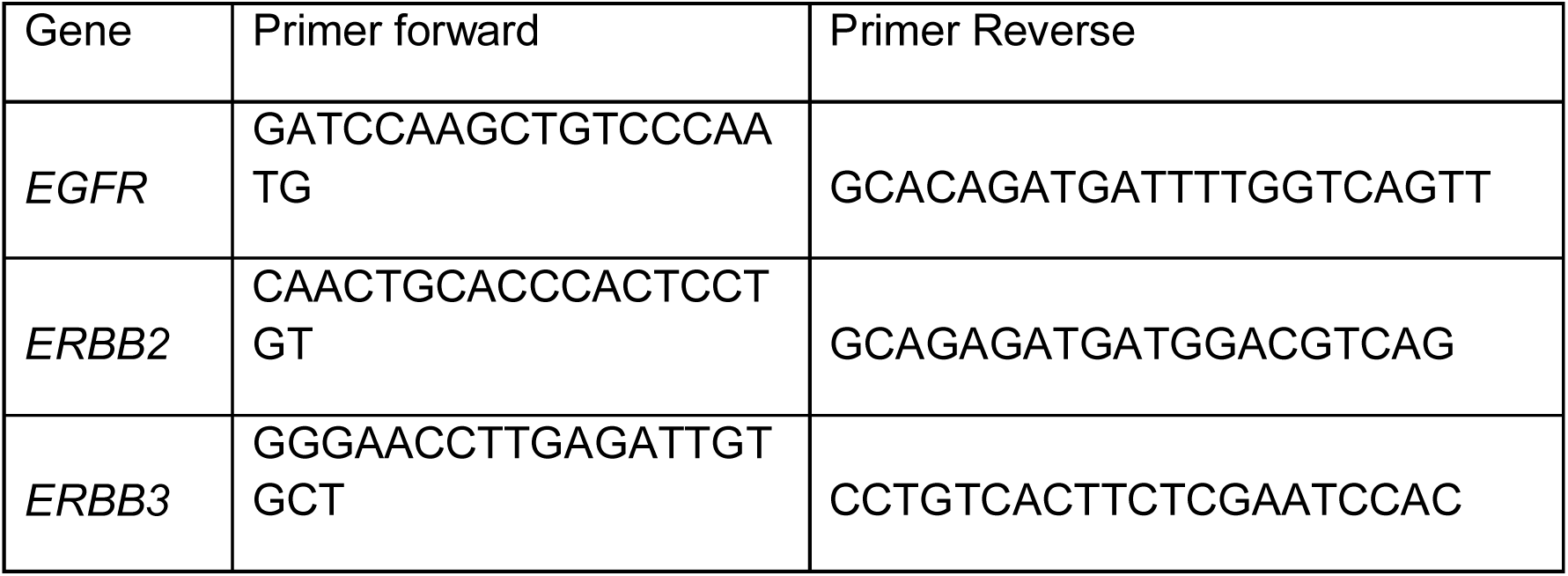

### Sanger sequencing

The coding exons and splice junctions of *PPARG* were amplified from genomic DNA by PCR with gene-specific primers available on request and sequenced by the Sanger method as described (22).

### WISP (Weighted In Silico Pathology)

WISP (https://cit-bioinfo.github.io/WISP/) is an approach to assess intra-tumoral heterogeneity from bulk molecular profiles. Based on predefined pure molecular or histological populations for a particular cancer type, this approach gives a fine description of each tumor in a standardized way. The methodology is based on non-negative least squares regression and quadratic programming optimization for estimating the mixed proportions of distinct populations for a tumor sample. It can be applied on transcriptomic or methylation data. The output is the mixing proportion estimations for all samples

For this analysis, we classified the samples from the MIBC CIT cohort (11) into luminal or basal subtype according to the BASE47 classifier (23) to refine pure samples and calculate pure population centroid profiles, using standard parameters. Then we estimated mixed proportions of pure populations for each our patient tumor and PDX samples, without scaling.

### Short tandem repeat signature

Patient tumors and corresponding PDX DNA samples were subjected to STR using the Authentifiler PCR amplification kit (Thermo Fisher Scientific, Illkirch, France) that amplifies 9 unique STR loci and the Amelogenin gender determining marker, according to the manufacturer instructions. PCR products were separated by capillary electrophoresis on ABI PRISM 3100 and results were analyzed using the GeneMapper software.

### Genomic alterations detection: mutations, copy number variant detection, variant calling and tumor mutational burden

Patient tumors and PDX were sequenced with a targeted NGS panel (called “DRAGON”) that has been developed by the genetics department of Institut Curie (Paris, France) and can detect mutations, copy number alterations (CNA), tumor mutational burden and microsatellite instability. It is composed of 571 genes of interest in oncology for diagnosis, prognosis and theragnostic (**Supplementary Table 1**). NGS primers were selected based on their specificity on the human genome. The whole method is described in **Supplemental Method Dragon**.

Deleterious genomic alterations were defined as follows: (i) for oncogenes, only gain-of-function mutations were considered (i.e., hotspots missense mutations, in-frame insertions/deletions/splicing described as oncogenic), (ii) for tumor suppressor genes, only loss-of-function mutations were considered (i.e., biallelic truncating alterations - nonsense mutations, frameshift insertions/deletions/splicing - or monoallelic truncating alterations associated with heterozygous deletion detected by copy number analysis).

### Gene expression analysis

#### Gene expression arrays/ Transcriptomic data/ Consensus Class

RNA of 31 PDX and patient tumors samples were hybridized in three batches in Affymetrix Human Genome U133 plus 2.0 Array Plates (Santa Clara, CA) according to Affymetrix standard protocols. Raw CEL files were RMA normalized (24) using R statistical software. PCA confirmed that no batch effect was observed. The arrays were mapped to genes with a Brainarray Custom CDF (Human EntrezG version 24) (25).

Molecular Consensus class were determined with the “consensusMIBC” R package (v1.1.0, https://github.com/cit-bioinfo/consensusMIBC) using the RMA-normalized transcriptomic data. For WISP, we used a previously published dataset (11), which contains Human MIBC samples (n = 85) also hybridized with Affymetrix Human Genome U133 plus 2.0 according to Affymetrix standard protocols. The Raw CEL files used here are available from ArrayExpress (http://www.ebi.ac.uk/arrayexpress/) under accession number E-MTAB-1803.

Raw CEL files were RMA-normalized using R statistical software. The arrays were mapped to genes with a Brainarray Custom CDF (Human EntrezG version 23) (25)

In both datasets, we obtained an log2-transformed expression matrix with one value per gene.

### Regulatory networks

The regulatory network was reverse engineered by ARACNe-AP (26) from human urothelial cancer tissue datasets profiled by RNA-Seq from TCGA (n=414). The RNA-seq data was downloaded from TCGA data portal using TCGAbiolinks package (R). Raw counts were normalized to account for different library sizes and the variance was stabilized with VST function in the DESeq2 R-package (27)

ARACNe was run with 100 bootstrap iterations using all probe-clusters mapping to a set of 1,740 transcription factors. Parameters used were standard parameters, with Mutual Information p-value threshold of 10−^8^.

### Regulon activity - VIPER

The VIPER algorithm tests for regulon enrichment based on gene expression signatures (28), using the regulatory network obtained from ARACNe on urothelial cancer, and we computed the enrichment of each regulon on the gene expression signature using different implementations of the analytic Rank-based Enrichment Analysis algorithm.

### Dual staining immunohistochemistry (IHC)

Dual immunostaining for GATA3 and KRT5/6 on FFPE samples was performed to screen for homogeneous or heterogeneous luminal or Ba/Sq tumors at the immunohistochemical level. Automated sequential dual staining immunohistochemistry (Discovery, Roche/Ventana, Tucson, AZ, USA) was used according to the manufacturer’s instructions. Tissue sections cut at 3 µm were dewaxed and subjected to antigen retrieval, then incubated first with a GATA3-specific rabbit monoclonal antibody (1:300, clone ZR65 Diagomics, Blagnac, France), followed by HRP-conjugated anti-rabbit IgG secondary antibody (MP-7401, Vector). The antigen–antibody reaction was detected using ImmPACT DAB reagent (SK-4105, Vector), producing brown staining in positive nuclei. In the second sequence, a primary rabbit monoclonal antibody against KRT5/6 (1:100, clone EP24/EP67; Diagomics, Blagnac, France) was incubated, followed by alkaline phosphatase-conjugated anti-rabbit IgG secondary antibody (ENZ-ACC110-0150, Enzo). The antigen–antibody reaction was revealed using ImmPACT red reagent (Vector), producing red staining in positive cytoplasm. Normal urothelium was used as a positive control. The staining for GATA3 and KRT5/6 was evaluated by one blinded pathologist (JF), providing respective quick scores (QS) calculated as intensity (0 to 3) multiplied by the percentage of stained tumor cells and normalized to [0;1]. Immunohistochemical thresholds for Ba/Sq tumors (IHC-Ba/Sq) defined as QS(KRT5/6) > 0.14 and QS(GATA3) < 0.02, or luminal QS(GATA3) >0.14 and QS(KRT5/6) < 0.02 were used (29). Tumors showing IHC-Ba/Sq and non-Ba/Sq areas were defined as having intra-tumoral heterogeneity.

### *In vivo* efficacy studies

Erlotinib (EGFR inhibitor) was purchased from MedChemExpress (Sollentuna, Sweden) and administered orally (gavage) five days per week during 4 weeks at a dose of 30 or 90 mg/kg (0.5% carboxymethylcellulose in PBS).

Erdafitinib (pan-FGFR inhibitor) was purchased from MedChemExpress (Sollentuna, Sweden) and administered orally (gavage) six days per week during 4 weeks at a dose of 10 or 30 mg/kg (20% 2-hydroxypropyl β-cyclodextrin in distilled water).

BGJ398 (pan-FGFR inhibitor) was purchased from LC Laboratories and administered orally (gavage) six days per week during 4 weeks at a dose of 30 mg/kg.

Cisplatin and Gemcitabine were purchased from Sigma (St. Quentin Fallavier, France). Both drugs were administered intraperitoneally at a dose of 60 mg/kg (NaCl 0.9%) once a week (D0, D7, D14) and 4 mg/kg (NaCl 0.9%) once every three weeks (D1), respectively.

For efficacy studies, mice were implanted as described above. Tumor fragments were transplanted into 6-week-old NMRI mice. When tumor reached a volume comprised between 60 and 270 mm^3^, mice were randomly assigned to the vehicle or treatment groups (n= 7-10). Tumor volume was calculated as: TV (mm3) = [length (mm) x width (mm)2]*π/6, where the length and width are the longest and shortest diameters of the tumor, respectively. Tumor volumes were then reported to the initial volume as relative tumor volume (RTV). Means of the RTV in the same treatment group were calculated. Growth curves were generated using the GraphPad Prism software.

### Western blot

Frozen PDX samples were resuspended in Laemmli lysis buffer [50 mM Tris-HCl (pH 6.8), 2 mM DTT, 2.5 mM EDTA, 2.5 mM EGTA, 2% SDS, 5% glycerol with protease inhibitors and phosphatase inhibitors (Roche)], and the resulting lysates were clarified by centrifugation. The protein concentration of the supernatants was determined with the BCA protein assay (Thermo Scientific, France). Proteins (10–50 μg) were resolved by SDS–PAGE in 10% polyacrylamide gels, electrotransferred onto Bio-Rad nitrocellulose membranes, and analyzed with antibodies against β-actin (Sigma Aldrich #A2228, used at 1/25,000), or the extracellular domain of FGFR3 (Abcam, # ab133644, 1/5,000). Anti-mouse IgG, HRP-linked, and anti-rabbit IgG, HRP-linked antibody (Cell Signaling Technology # 7076 and # 7074, used at 1/3,000, Saint-Cyr-L’École, France) were used as secondary antibodies. Protein loading was checked by Amido Black staining of the membrane after electrotransfer.

### Statistical and bioinformatic analysis

Comparisons between PDX treatment responses were done using Mann-Whitney test.

Bioinformatics analyses was performed with R (4.0.2).

## Results

### Establishment of urothelial PDXs

We successfully obtained 31 PDXs from 153 tumors (global engraftment rate ∼20%, median latency 27.5 days with a range of 10-70 days) from January 2009 to October 2019. The main clinico-pathological characteristics of the patients and patient tumors are summarized in **Table 1** (additional clinical data from patients are presented in **Supplementary Table 2**). As expected, the majority of PDXs was derived from male patients (84%), with bladder as the primary site (84%) and were urothelial carcinomas (90%). The frequency of squamous cell carcinomas (SCC, 3/31, 10%) was higher than the known frequency of these tumors, potentially owing to a higher engraftment success rate (n=4/10, with one of the 4 that could not be further analyzed due to patient serology). One PDX was derived from a non-muscle-invasive bladder cancer (NMIBC). Four patients had received systemic or radiotherapy treatment before the establishment of the PDX.

**Table 1.**
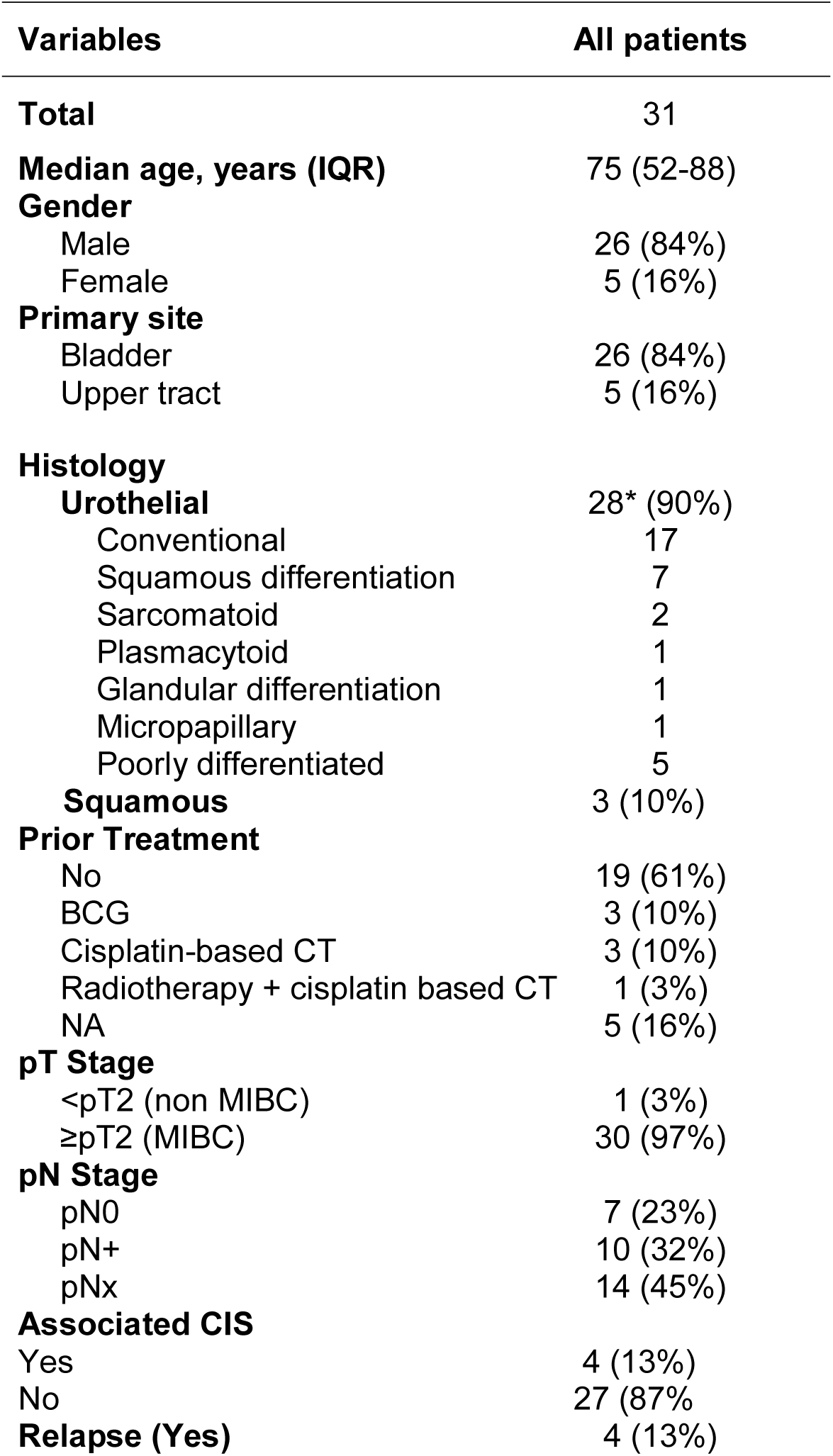
Clinico-pathological characteristics of the samples used for PDX. CT: Chemotherapy * Variant total is above n=31 as some tumors have multiple variants

### Histological and genomic characterization of PDXs

Histopathological analysis of H&E-stained slides of patient tumors and PDXs was performed by two pathologists (VL, JF), based on the current WHO Classification of Tumors of the Urinary System (30). High histological concordance between patient tumors and PDXs was observed (**FIG1A**) with the exception of 3 PDXs showing distinct histology compared to the matched patient tumors (M1030 and BLCU-011 lost the variant observed in the tumor, while more squamous variant was observed in R1056, **Supplementary Table 2**). We confirmed by short tandem repeat (STR) profiling the concordant genetic identity between patient tumor and derived PDXs (**Supplementary Table 3)**, with 85 to 100% of conserved STR for the different models, as example for L987 in **FIG1B**. We characterized genomic alterations for 571 cancer-related genes (**Supplementary Table 1 and 4**) in all 31 PDXs using a targeted next-generation sequencing assay allowing the detection of mutations, estimation of copy number alterations (CNA), tumor mutational burden (TMB) and MSI status (**FIG1C**). As anticipated, the most frequent genomic alterations were mutations in *TERT* (68%) and *TP53* (61%) and homozygous deletion of *CDKN2A/B* (∼50%). We also identified activating mutations in potentially actionable genes such as *PIK3CA* (19%), *ERBB2* (19%), *FGFR3* (13%), *BRAF* (6%), *ERBB3* (3%), *KRAS G12C* (3%), and truncating mutations in the epigenetic genes *KDM6A* (19%), *ARID1A* (23%), *KMT2D (*19%), *KMT2A/B/C* (3% each) and *ARID2* (3%). One UTUC PDX displayed microsatellite instability associated with a bi-allelic deletion of *MSH2* and this alteration was also observed in the parental tumor. To determine whether the PDXs retain the genomic alterations of the matched patient tumor, we sequenced 5 patient tumor/PDX pairs. The overall concordance of observed genomic alterations was high (90-100%) (**FIG1D**), except for the MSI-High PDX (B521) that harboured high TMB (40%). PPARG pathway activation, through PPARG amplifications or RXRA and PPARG activating mutations, is a known key feature of luminal tumors and a potential therapeutic target (12,22,31–33). We did not observe any *RXRA* mutation in our sequencing analyses. Since *PPARG* was not in our targeted panel, we performed Sanger sequencing of the hotspot region within the ligand binding domain of *PPARG* (22) and identified one patient tumor/PDX pair (B521) harbouring the T475M activating mutation (n=1/23 tested PDX) and one patient tumor/PDX pair (M559) harbouring the non-characterized and non-recurrent L339F mutation (22).

**Figure 1.**
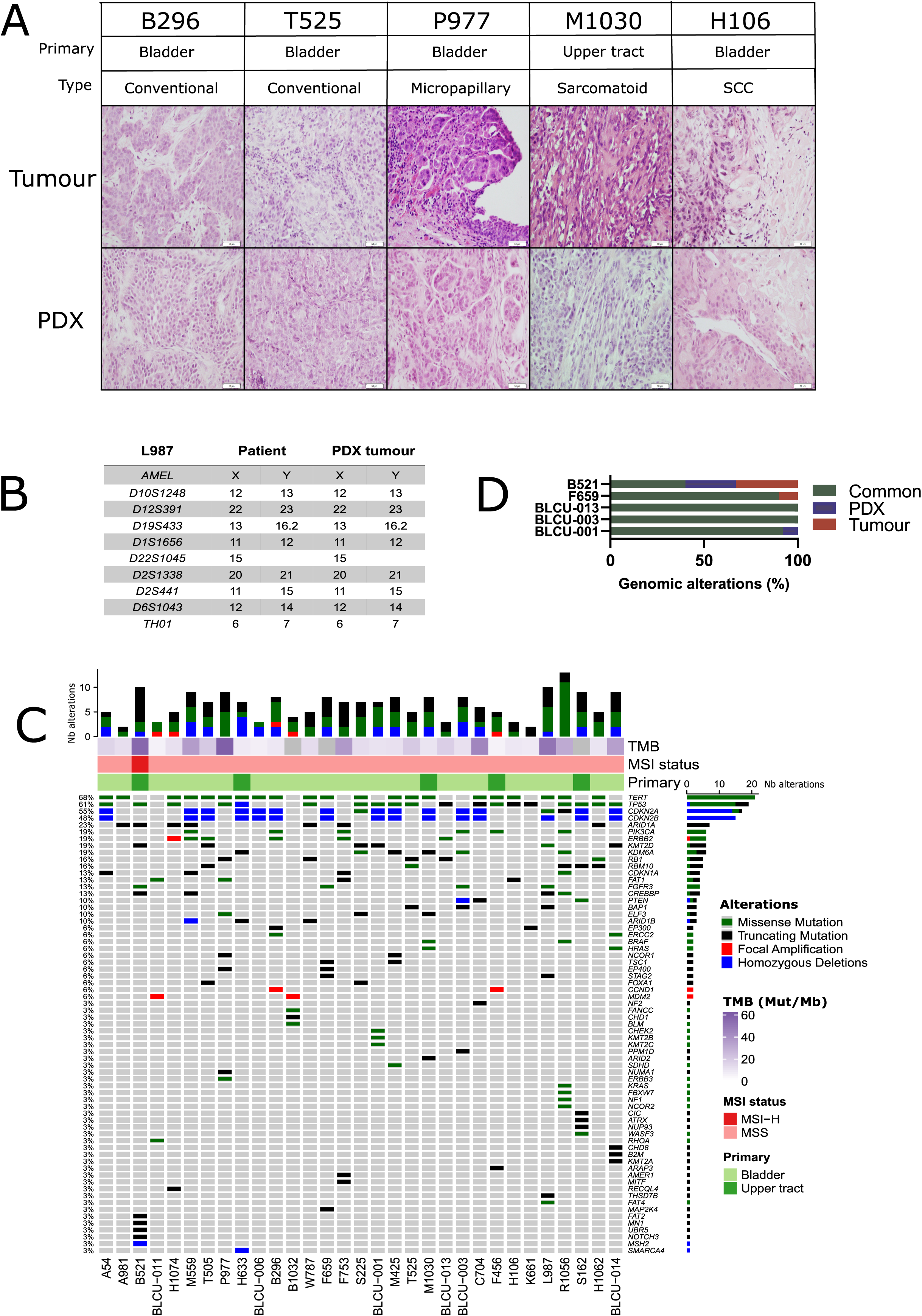
Histological and genomic characteristics of patient tumors and paired derived PDXs. **A-** Histology of bladder/ureteral patient tumors and corresponding PDX models demonstrated similar histological features as assessed by H&E staining. Scale bar corresponds to 50□µm. H&E slides of a section of each patient tumor and matching PDX were reviewed by a board-certified pathologist and representative pictures for the different histologies or variants are shown. **B-** Short tandem repeat (STR) signature of patient specimen and PDX tumor, example of the L987 case. **C-** Somatic genomic landscape of 31 bladder and ureteral PDXs analyzed using an in-house targeted sequencing assay (571 cancer-related genes, supplementary Table 2-3), tumor mutational burden per megabase (TMB, indicated in log2 scale for each sample) and microsatellite instability-high (MSI-H) versus microsatellite stable (MSS) status. **D-** Concordance of genomic alterations in 5 pairs tumor/PDX

### PDX recapitulate the molecular subtypes and intra-tumoral heterogeneity

We then sought to determine whether PDX models recapitulate the molecular subtypes of patient tumors. In total, transcriptomic data using Affymetrix U133plus2 array were available for 22 patient tumor/PDX pairs and 8 individual PDXs. Unsupervised clustering analysis using the top 200 most variant genes did not highlight any segregation in PDXs by primary site or histological tumor (**Supplementary Figure 1,** with exclusion of the NMIBC PDX for the transcriptomic subtype analysis). We therefore considered all patient tumors/PDX similarly and stratified them into 6 subtypes by applying the molecular consensus classifier (5) (**FIG2A**). We observed 15 Ba/Sq (52%), 11 luminal-papillary (38%) (LumP) and 3 luminal-unstable (LumU) (10%) PDXs. All 3 SCC patient tumors/PDXs classified as Ba/Sq. In contrast to genomic and histological characteristics, transcriptomic profiles were less stable between patient tumor/PDX pairs. Specifically, 8 patient tumors gave rise to a PDX with a distinct molecular subtype (36%) including 5 Ba/Sq patient tumor to LumP PDX, one LumP to LumU, one Stroma-rich to LumU and one LumNS to LumP subtype. We also stratified patient tumors and PDXs according to the BASE47 classifier (23) (**FIG2A**). Among the 16/22 patient tumors that classified as basal, 6 gave rise to a luminal PDX. In contrast, all luminal tumors formed luminal PDXs. For 6 matched tumor-PDX pairs, for which we did not have transcriptomic data, we performed GATA3 (luminal) and KRT5/6 (basal) dual immunostaining (**FIG3C**) to assign molecular subtypes using previously defined thresholds for each marker (29) and we did not observe a difference in subtype between tumors and PDXs (**FIG2A**). To explore whether the basal to luminal discordance could be related to intra-tumoral heterogeneity, we evaluated the molecular heterogeneity in both patient tumors and PDXs using WISP algorithm (Weighted In Silico Pathology) (**FIG2A**). We observed admixed proportions of luminal and basal subtypes in 59.1% of patient tumors and 41.4% of PDXs, including all 6 cases with discordant BASE47 subtypes. The high molecular heterogeneity found in the 6 basal tumors that gave rise to luminal PDXs was conserved in most of the matched PDXs. These findings suggest an intrinsic plasticity of these tumors leading to a shift in subtype, rather than a sampling bias of an area of a given subtype within a molecularly heterogeneous patient tumor (**FIG2A**, lower panels).

**Figure 2:**
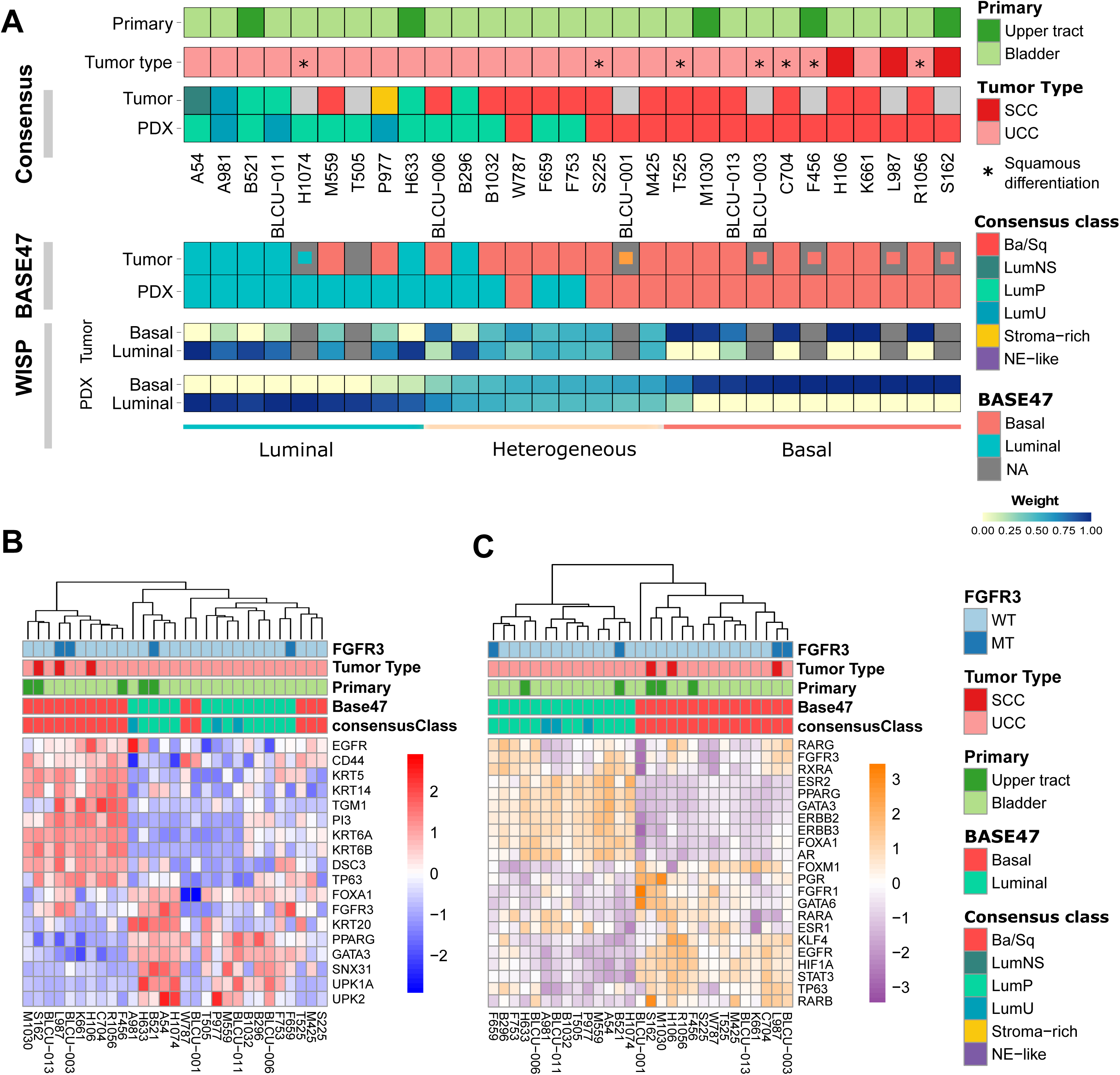
Transcriptomic analysis of patient tumors and paired PDXs. **A-** Tumors and PDXs were classified into 6 subtypes using transcriptomic Affymetrix U133plus2 array data according to the molecular consensus classifier developed for MIBC, corresponding box colors indicated in legend on the right. 22 patient tumor-PDX pairs and 7 individual PDXs were analyzed, where tumors without transcriptomic data are indicated in grey. Urothelial carcinomas with divergent squamous differentiation are highlighted with *.**B-** Upper panels: Tumors and PDXs were classified using transcriptomic data as luminal and basal subtypes according to the BASE47 classifier (23). In patient tumors with missing transcriptomic data, we assessed the luminal (blue), basal (red) or heterogeneous (orange) subtype by immunohistochemistry (inset small boxes), as defined in methods. Lower panels: intra-tumoral heterogeneity and proportion of luminal and basal subtype admixture as evaluated from transcriptomic profiles using WISP algorithm (Weighted In Silico Pathology). Based on the PDX WISP results, samples were molecularly classified as luminal, basal or heterogeneous. **B-** Heatmap of PDX samples based on gene expression of selected luminal or basal markers. **C-** Heatmap of PDX samples based on the regulon activity of the main regulators previously identified within the different molecular subtypes of MIBC (5,6)

**Figure 3:**
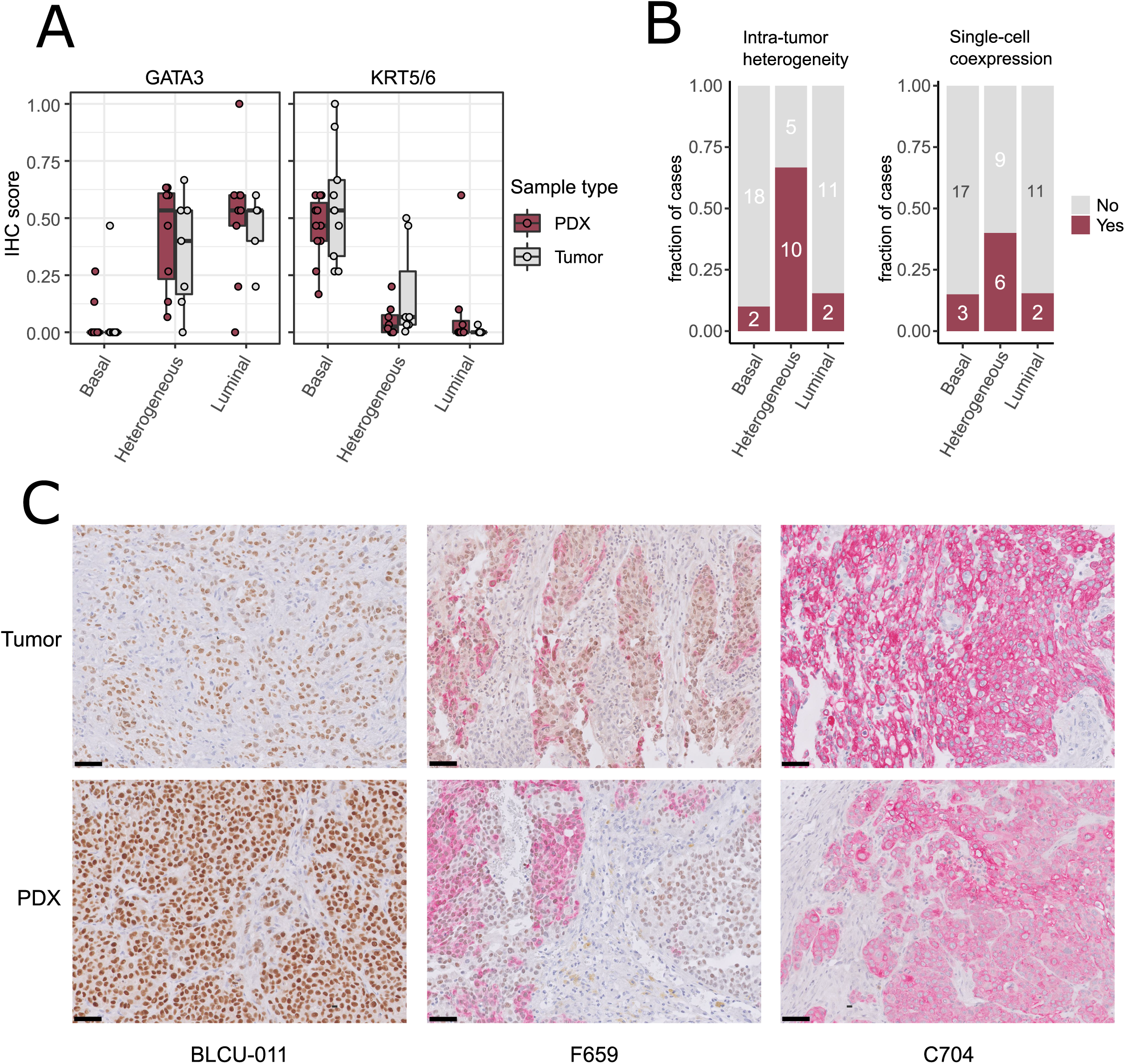
Intra-tumor heterogeneity in tumors and PDXs at the protein level by immunohistochemistry. A- GATA3 and KRT5/6 expression levels (normalized quick scores), grouped according to the PDX WISP molecular classification (luminal, heterogeneous and Ba/Sq, FIG 2A). B- Proportion of tumors with intra-tumoral heterogeneity (left) and GATA3 + KRT5/6 co-expression at the single cell level (right), grouped according to the PDX WISP molecular classification (luminal, heterogeneous and Ba/Sq, FIG 2A). C- Patterns of dual immunohistochemistry staining for GATA3 (brown, nuclear) and KRT5/6 (red, cytoplasmic) in paired tumor/PDX of a luminal (BLCU-011), a heterogeneous (F659) and a Ba/Sq (C704) example.

Using defined biomarkers of luminal and basal differentiation (5,6), we confirmed that PDXs were globally separated between luminal, differentiated tumors and basal tumors (**FIG2B** **and Supplementary FIG1**). To further explore the transcriptomes of our PDXs, we inferred the activity of 22 major regulons, as defined in TCGA analysis (6) (**FIG2C**). Using this approach, we observed a perfect separation of luminal and basal PDXs in clustering analysis, independently of the primary site of patient tumor. Similar to patient tumors from TCGA, we observed a higher *EGFR* regulon activity in basal PDXs and higher *PPARG/ERBB2/ERBB3* regulon activity in luminal PDXs (6). Of note, *FGFR3* mutations are enriched in LumP tumors (5) and we observed *FGFR3* mutations in 2 LumP PDXs, including 1 derived from UTUC, but also in 2 Ba/Sq PDXs, including 1 derived from a SCC tumor (**FIG2B** **and** **FIG2C**) (5). As expected, high FGFR3 regulon activity was observed in tumors bearing *FGFR3* mutations.

To validate the intra-tumoral molecular subtype admixture inferred from the transcriptomic data *in situ*, we performed dual immunohistochemistry staining combining a luminal (GATA3) and a basal marker (KRT5/6) in a subset of patient tumors and PDXs. We confirmed the presence of subtype marker expression admixture *in situ*, with either spatially distinct areas showing different IHC profiles (defined hereafter as intra-tumoral heterogeneity), or single-cell coexpression of the two markers, or a combination of both patterns of admixture. By comparing the results between tumors/PDXs classified as basal, luminal or heterogenous based on the WISP analysis (**FIG2A**, **FIG3**), we observed that tumors/PDXs classified as pure luminal or Ba/Sq based on transcriptomic data displayed higher GATA3 or KRT5/6 staining levels, respectively (**FIG3A**). The WISP heterogeneous samples had more intermediate staining levels of both GATA3 and KRT5/6 (**FIG3A**) and showed more intra-tumoral heterogeneity compared to pure Ba/Sq or luminal cases (10/15 vs 4/29, p<0.01) (**FIG3B**). Among samples with intra-tumoral heterogeneity, a GATA3-KRT5/6 co-expression at the single cell level was also identified in 9/14 samples (**FIG3B-C**).

### Chemosensitivity of PDXs

Cisplatin-based chemotherapy being the standard of care of MIBC and high risk UTUC, we assessed the sensitivity of 10 PDXs representative of the different subtypes (6 basal, 4 luminal) to cisplatin plus gemcitabine (**FIG4**). We did not observe a significant difference between the proportion of basal (5/6, 83%) and luminal PDXs (2/4, 50%) with significant growth inhibition upon treatment (Fisher’s Exact Test p=0.5). Due to the low number of recurrent genomic alterations and the number of models tested, it was not possible to statistically explore the association between genomic alterations and chemosensitivity.

**Figure 4:**
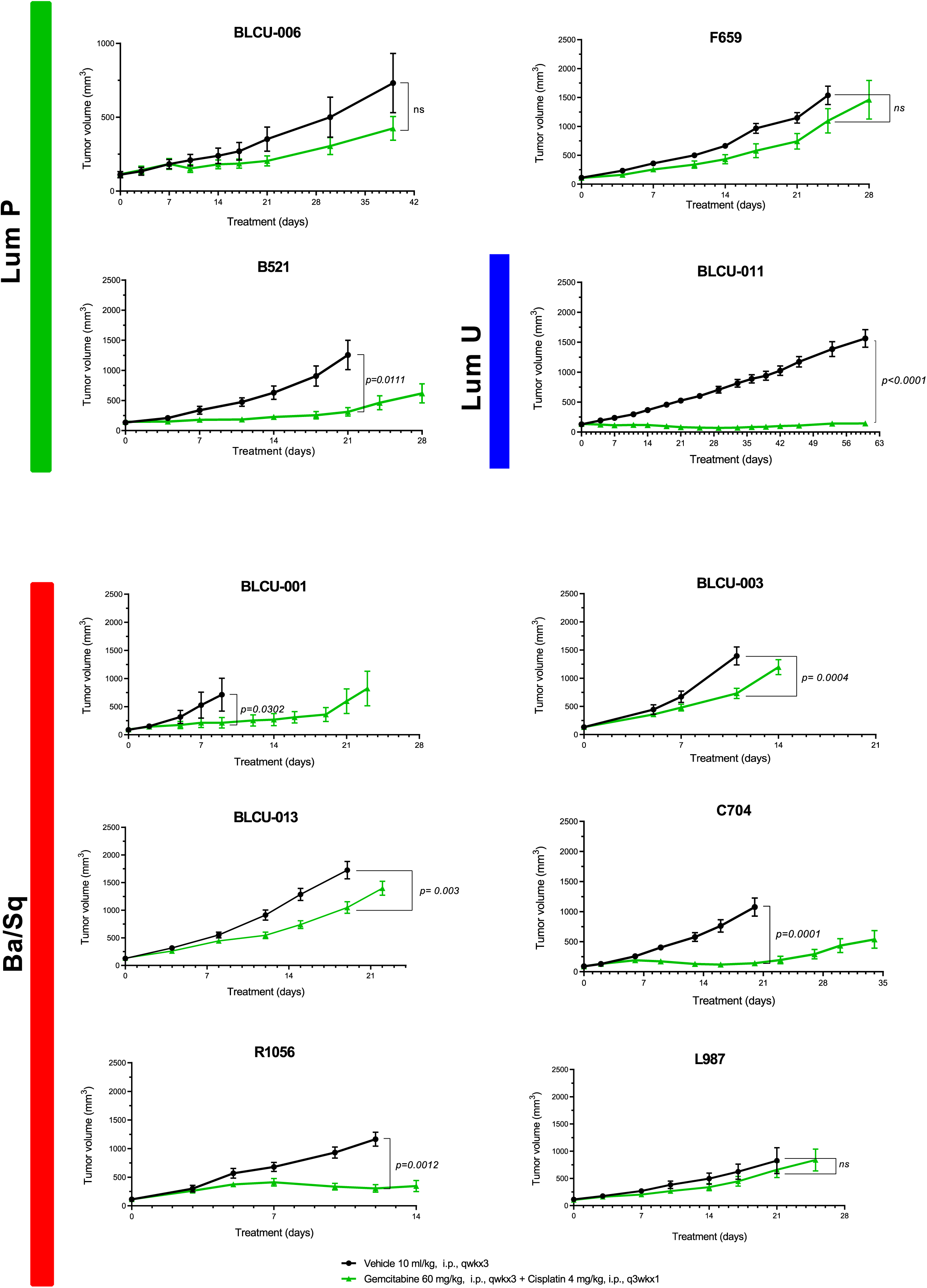
Chemosensitivity of representative basal and luminal PDX models. Mice with established PDXs (67-270 mm^3^) were treated with cisplatin plus gemcitabine (green) as indicated and control mice were treated with vehicle alone (black) (n = 7 to 10 animals per group). Tumor size was measured at the indicated time points. Data are presented as mean ± SEM. Results were compared using Mann-Whitney test.

### EGFR targeted therapy is effective in non-sarcomatoid Ba/Sq PDXs

In relation to the high EGFR activity in basal tumors, we have previously shown that EGFR is an effective therapeutic target in different *in vivo* basal preclinical models (**11**). The aggressive sarcomatoid variant of MIBC is suggested to occur through progression of Ba/Sq tumors (34). We recently analyzed sarcomatoid tumor transcriptomes and identified a loss of EGFR regulon activity during progression of Ba/Sq tumors to the sarcomatoid variant (Fontugne *et al.,* unpublished results). In line with these findings, we identified a low EGFR regulon activity in a PDX (BLCU-001) compared to the other Ba/Sq PDXs, which was classified as a Ba/Sq sarcomatoid tumor (**FIG2C** and **supplementary Table 1**). To assess whether the loss of EGFR activity could impact the sensitivity to EGFR inhibition, we compared the effect of anti-EGFR treatment (erlotinib) in two Ba/Sq models presenting high EGFR-regulon activity (L987 and H106, **FIG2C**) and BLCU-001 (**FIG5A**). As expected, the two SCC Ba/Sq models were sensitive to Erlotinib whereas the Ba/Sq model with sarcomatoid differentiation was resistant (**FIG5A**). Of note, two other Ba/Sq PDXs (C704 and R1056) were sensitive even when we used a lower dose of erdafitinib in a second experiment (**FIG5B**).

**Figure 5:**
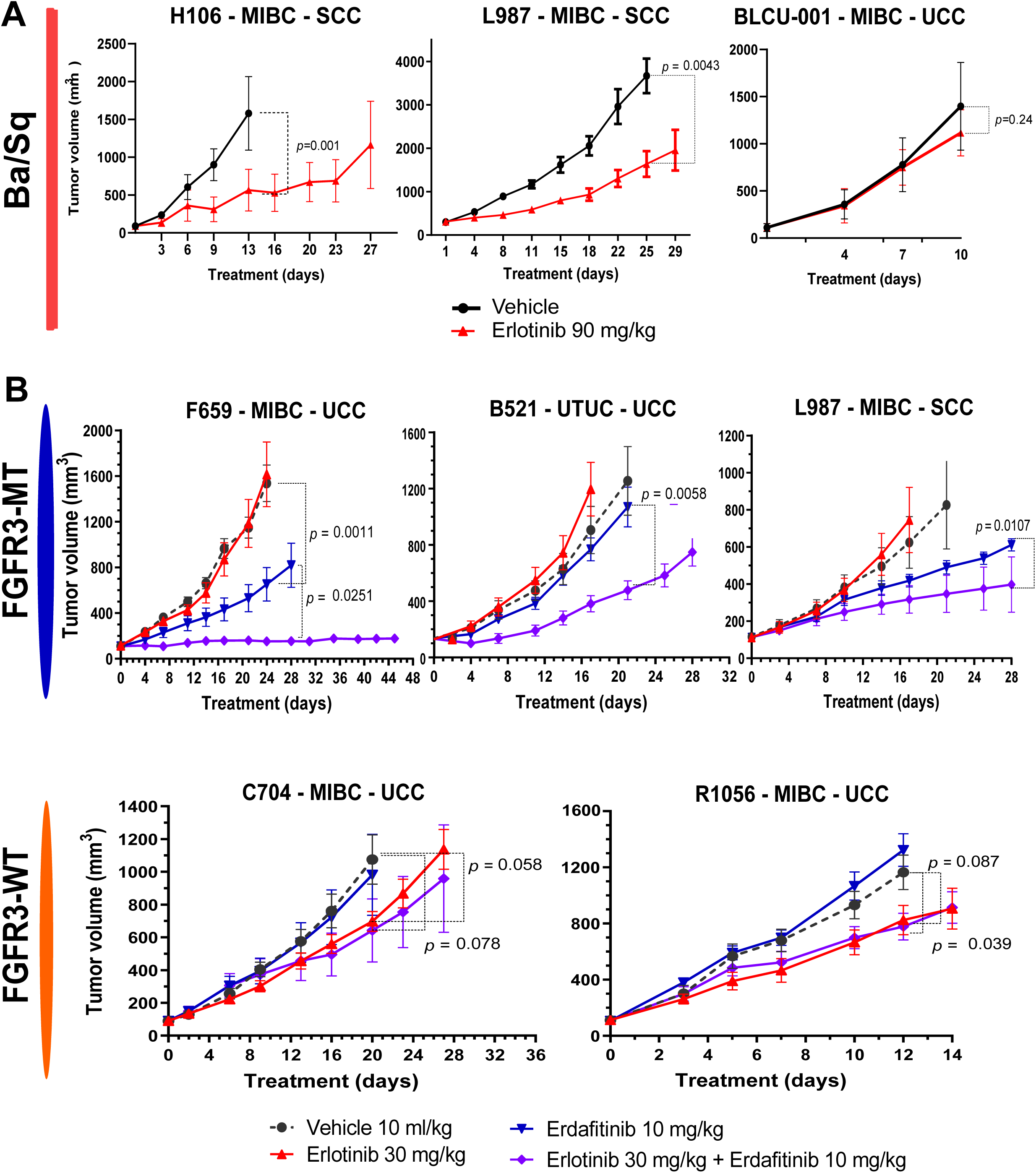
Sensitivity to anti-EGFR or the combination of EGFR and FGFR inhibitors in Ba/Sq and *FGFR3*-mutated PDXs. A-Mice with established basal/squamous (Ba/Sq) PDXs (67-270 mm3) were treated with anti-EGFR (erlotinib 90 mg/kg, red) or control vehicle alone (black). B-Mice with established *FGFR3*-mutated tumors and controls were treated with control vehicle (black), low dose anti-EGFR (erlotinib 30mg/kg, red), pan-FGFR inhibitor (erdafitinib 10mg/kg, blue), or the combination (purple), as indicated (n = 7 to 10 animals per group). Tumor size was measured at the indicated time points. Data are presented as mean ± SEM. Results were compared using Mann-Whitney test.

### Combined inhibition of FGFR and EGFR improves response of *FGFR3*-mutated PDX compared to FGFR inhibition alone

FGFR inhibitors have already demonstrated clinical efficacy in FGFR3-altered tumors (4). However, resistance to treatment is systematically observed over time. Consistently, treatment of two FGFR3-mutated PDXs, one Ba/Sq and one LumP, both presenting high expression levels of FGFR3 protein compared to other non-mutated PDXs, with a pan-FGFR-inhibitor (BGJ398) (**Supplementary FIG2A**) only reduced tumor growth (**Supplementary FIG2B**). Different pre-clinical studies using bladder-derived cell lines identified EGFR activation as a mechanism of resistance to FGFR inhibition (35–37). Using RT-qPCR, we confirmed that this mechanism could be relevant in our PDX models since anti-FGFR treatment induced the overexpression of EGFR but also ERBB2 and ERBB3 in our *FGFR3*-mutated L987 model (**Supplementary FIG2C**).

Therefore we then tested if a combination of FGFR3 and EGFR/ERBB2 inhibition could overcome the compensatory upregulation of EGFR signalling and increase the sensitivity of FGFR3-mutated PDXs to FGFR inhibition, as previously observed for RT112 xenografts harbouring FGFR3-TACC3 fusion (35). We treated 5 PDXs (3 *FGFR3*-mutated: L987, B521, F659; 2 Ba/Sq *FGFR3*-WT: R1056, C704) with a pan-FGFR inhibitor (erdafitinib) that recently obtained FDA approval for locally advanced or metastatic MIBC with *FGFR3* genomic alterations and an EGFR inhibitor (erlotinib, which has been described to have ERBB2 inhibitory effect (38)) alone and in combination (**FIG5B**). To limit toxicity, we used suboptimal doses for each inhibitor when administered in combination. In these conditions, monotherapies presented limited effects, with significant growth inhibition in two *FGFR3*-mutated and in two basal PDX upon erdafitinib and erlotinib treatment, respectively. The combination of erlotinib plus erdafitinib led to an improved response in all *FGFR3*-mutated PDXs compared to each of the monotherapies, while no additional benefit was observed for the combination in the Ba/Sq *FGFR3*-WT PDXs (**FIG5B**).

## Discussion

The development of relevant preclinical models closely mimicking patient tumors is key in the identification of new potential therapies for precision medicine and PDXs are widely used in this regard in oncology (20,39).

We report here, like others (18–21), that PDXs derived from bladder and ureteral cancers preserved histological and genomic properties of patient tumors. However, their transcriptomic profiles were less stable. Namely, we identified basal patient tumors giving rise to luminal PDXs. For these cases, both tumors and PDXs revealed high heterogeneity, suggesting an intrinsic cell plasticity. Such heterogeneity and cell lineage plasticity were recently demonstrated by Sfakianos et al. in the basal N-butyl-N-(4-hydroxybutyl)-nitrosamine (BBN) chemically-induced mouse bladder tumors using both single cell transcriptomic analysis and FACS analysis after in vivo transplantation of FACS pre-sorted and cultured tumor cells (40). Basal tumors present a more abundant stroma compared to luminal tumors including high infiltration by immune cells (5). Our results also suggest that, because the stroma in immuno-deficient mice differs from human tumor stroma, the cross-talk between tumor and stroma may impact tumor cell phenotype. Such cross-talk has been reported in 3D-*ex vivo* models, where the absence of cancer-associated fibroblasts induced a shift from luminal tumors to basal organoids (41,42) whereas basal organoids engrafted in mice then developed a luminal phenotype (41). Interestingly, the molecular heterogeneity of luminal/basal markers observed in tumor and PDX pairs also reinforces the relevance of our models in drug efficacy evaluation, given that tumor heterogeneity is a known cause of resistance to treatment (43).

We described potentiall actionable genomic alterations in our PDXs, such as *PIK3CA* mutation, *ERBB2* mutation/amplification or *MDM2*-amplification (44,45), whose frequencies are comparable with TCGA (6). Epigenetic drugs could also represent a promising therapeutic approach in monotherapy or in combination when considering the high proportions of PDXs harbouring at least one mutation in epigenetic genes (46). Our models could thus be useful to evaluate such new potential therapies.

Previously, we proposed EGFR as a therapeutic target in basal MIBC based on findings from different *in vitro* and *in vivo* preclinical models (11). Here, we further validated the EGFR dependency of Ba/Sq tumors using PDX models. However, anti-EGFR treatments have shown disappointing results in the clinic, though some pathological responses have been observed in the neoadjuvant setting (47). This discrepancy between preclinical and clinical settings warrants further investigation. We hypothesize that the rich stroma of basal tumors contributes to anti-EGFR resistance, as recently demonstrated with cancer-associated fibroblasts in lung cancer (48). Indeed, rich stroma is absent in our preclinical models, with the exception of the BBN model, which presented only a moderate sensitivity to anti-EGFR treatment (11). Our results also suggest that sarcomatoid differentiation could be another mechanism of resistance to anti-EGFR treatment. In agreement, sarcomatoid differentiation is linked to EMT, which was previously shown to impair anti-EGFR sensitivity in lung cancer (49).

In agreement with previous studies (50,51), we found that upper urinary tract and bladder urothelial carcinomas harboured similar genomic alterations but at different frequencies, except for the presence of MSI-H in one UTUC (50,51). The diversity of our bank—including both MIBC (Urothelial (UCC) and Squamous (SCC)) and UTUC—allowed us to observe that urothelial carcinoma PDXs originating from the upper urinary tract or the bladder were also highly similar at the transcriptomic level. Additionally, we found that SCC samples did not classify into a molecularly distinct group of tumors, instead grouping with urothelial Ba/Sq carcinomas. Whereas up to now SCC, UCC and UTUC were considered as highly different entities in terms of response to treatment, we found that the same molecular classifications/ genetic alterations can predict the same therapeutic response for the three entities. Indeed independently of cell of origin, basal PDXs were sensitive to anti-EGFR therapies, except for sarcomatoid tumors, which are characterized by low EGFR activity. Furthermore, most *FGFR3*-mutated PDXs were sensitive to FGFR inhibition independently of their cell of origin or their luminal/basal phenotype.

Finally, it has been shown in RT112 xenografts (bladder cancer cell line presenting an FGFR3-TACC3 fusion observed in 3% of MIBC) that a combination of anti-EGFR/anti-FGFR was more potent than anti-FGFR alone (35). We validated here these results using our more clinically relevant PDX models with FGFR3 mutations (observed in 15% of MIBC) (3 different models derived from UTUC, MIBC-UCC and MIBC-SCC) reinforcing the interest to further test this combination treatment in the clinics. For treatment optimization and to limit side effects and toxicity, it will be of interest to explore if this effect is more specifically associated with EGFR or HER2 inhibition.

## Conclusion

Muscle-invasive bladder and upper urinary tract cancers are heterogeneous and aggressive diseases with no satisfactory treatment. We have developed and characterized highly relevant preclinical models for bladder and upper tract carcinoma, recapitulating the molecular heterogeneity and drug responses observed in the clinic. Our work supports a benefit of combined FGFR and EGFR inhibition in FGFR3-mutated tumors. Overall, our models represent an essential tool for the development of new efficient therapies against these aggressive cancer types.

## Supporting information

Supplemental Method Dragon

## Availability of data and materials

All data requests should be submitted to the corresponding authors for consideration. Access to anonymised data may be granted following review.

Different modalities of access to PDX models will be possible though Urosphere company (commercially available, outsourcing or collaboration) and request should be submitted to the corresponding authors.

Transcriptomic data of PDXs and tumors are available in GEO database (GSE181962).

## Acknowledgments

We thank David Gentien from the genomics platform of Institut Curie.

We also thank the Alsace incubator SEMIA, Strasbourg, France, for advices and financial supports (Recipients TM and HL for Urolead SAS). We also thank the competitivity pole Alsace Biovalley, Illkirch-Graffenstaden, France for advices and labelization (TM and HL, for Urolead SAS). Finally, we also thank Alsace Satt Conectus, Illkirch-Graffenstaden, France, for technologies transfer and advices (TM and HL, for Urolead SAS).

## Author’s contribution

Isabelle Bernard-Pierrot had full access to all the data in the study and takes responsibility for the integrity of the data and the accuracy of the data analysis.

HL, CB, ML and LC designed, performed experiments or bioinformatics analyses, analyzed and interpreted the data. HL, CB and ML developed the PDXs models. HL, XG, PR provided samples and helped with clinical data analysis. CB and ML performed the pharmacological characterization of the PDXs. CB, CK and JF prepared DNA and RNA for genomics analysis. CB performed STR analysis under TM supervision. CK performed and analyzed sanger sequencing. CK and VD performed immunohistochemical experiments under YA supervision. JF performed histo-and immunohisto-pathological analysis and designed pharmacological studies for sarcomatoid tumors under YA supervision. FD performed Western-blot and RT-qPCR analysis. LC performed most of the bioinformatics analyses together with help of CSG, XM and AK for tumor’s/ PDXs’ classification and for regulon analysis. ADR supervised AK and CSG. EC normalized and annotated Affymetrix array data and centralized the data for bioinformatics analyses. JMP and RB performed and analyzed targeted NGS under the supervision of IB. VL performed and supervised histopathological analysis of tumors and PDXs. YM and PL designed the study and supervised establishment of part of the PDXs and pharmacological characterization of PDXs. FR supervised the genomic analyses. TM and HL designed and supervised the establishment of part of the PDXs. IBP and TM designed and supervised the research, analyzed and interpreted the data. LC, CB, JF, TM and IBP wrote the paper. All authors critically read and contributed to the final version of the manuscript.

## Ethics Declaration

### Ethics approval and consent to participate

All patients provided written informed consent. All research in this study conformed to the principles of the Helsinki Declaration.

### Consent for publication

All patients consent to participate

## Abbreviations

PDX: Patient derived xenografts
NMIBC: non-muscle-invasive bladder cancer
MIBC: muscle-invasive bladder cancer
FGFR: fibroblast growth factor receptor
EGFR: epidermal growth factor
SCC: squamous cell carcinoma
UCC: urothelial cell carcinoma
UTUC: upper urinary tract urothelial carcinomas

## Figure legends

**Supplemental Figure 1: Unsupervised hierarchical clustering and heatmap of PDX samples based on genes with the most variant expression (n=200)**

PDX identifiers indicated at the bottom. Classifications indicated according to legend on the right (as in Fig 2).

**Supplemental Figure 2: FGFR3 expression in PDXs and sensitivity to anti-FGFR of FGFR3 mutated PDXs**

**(A)** FGFR3 protein levels (Western blot) of the different PDX models. The urinary bladder cancer cell line, RT112, was used as control for WT FGFR3 (black arrow) and FGFR3-TACC3 (white arrow) protein expression. Beta-actin was used as loading control. Asterisk indicates non-specific band.

**(B)** Mice with established *FGFR3*-mutated (FGFR3-MT) PDXs (67-270 mm3) were treated with a pan-FGFR inhibitor (BGJ398). Control mice were treated with vehicle alone (n = 7 to 10 animals per group). Tumor size was measured at the indicated time points. Data are presented as mean ± SEM. Results were compared using Mann-Whitney test.

**(C)** EGFR, ERBB2 and ERBB3 expression (RT-qPCR analysis) in L987 model at the end of treatment in mice treated with vehicle or with the pan-FGFR inhibitor BGJ398.

**Supplementary Table 1:** Clinical and histological characteristics of patient tumors and matched PDXs.

**Supplementary Table 2:** Short Tandem repeat (STR) profiling of patient tumors and matched PDX(s)

**Supplementary Table 3:** List of the sequenced genes in the next-generation targeted sequencing panel.

**Supplementary Table 4:** Next-generation sequencing results (mutations, copy number alterations, microsatellite status and tumor mutational burden).

