## Supplementary figures and images for "Integrated molecular and pharmacological characterization of patient-derived xenografts from bladder and ureteral cancers identifies new potential therapies"

### Supplemental Figure 1

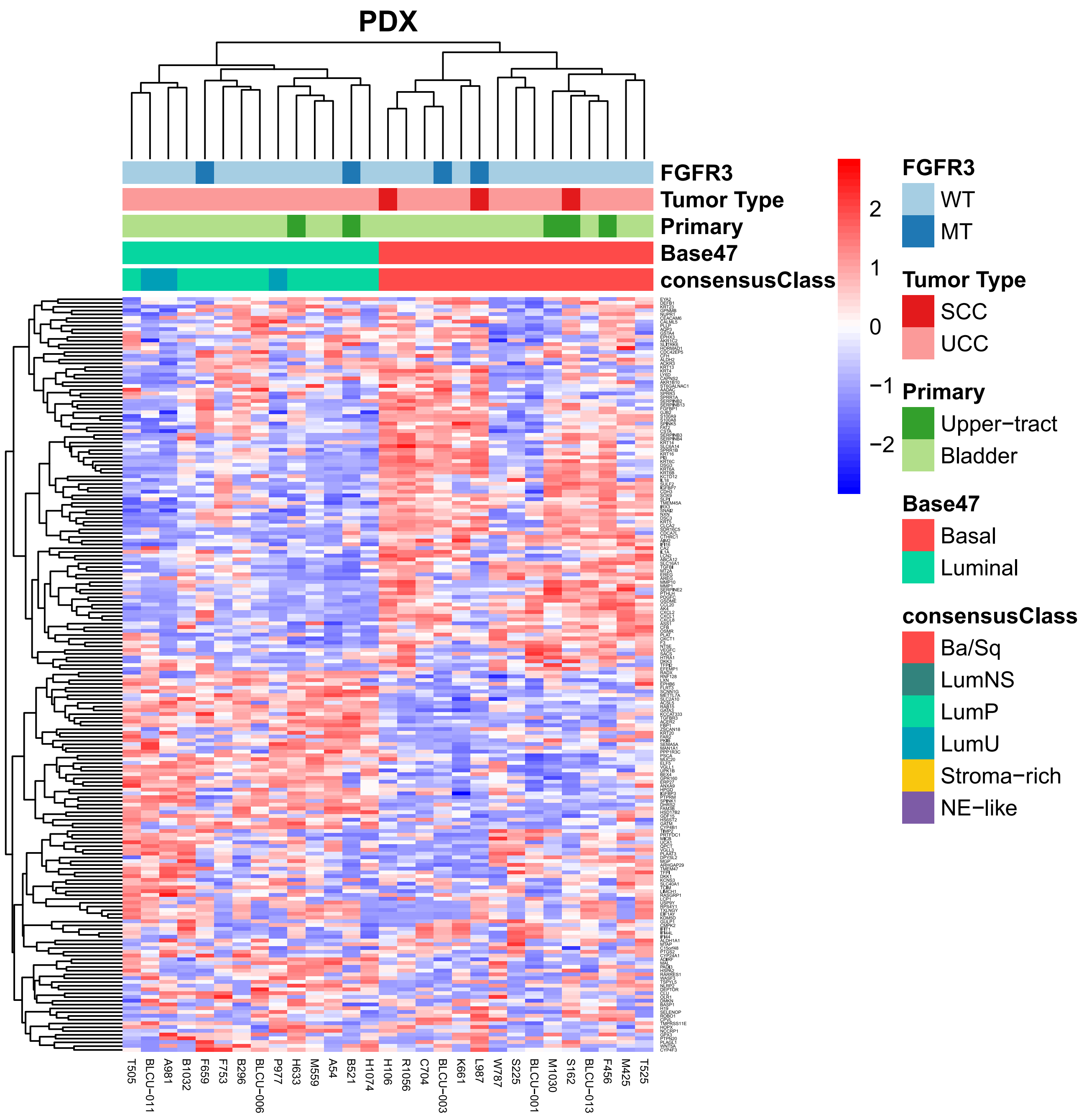

### Supplemental Figure 2

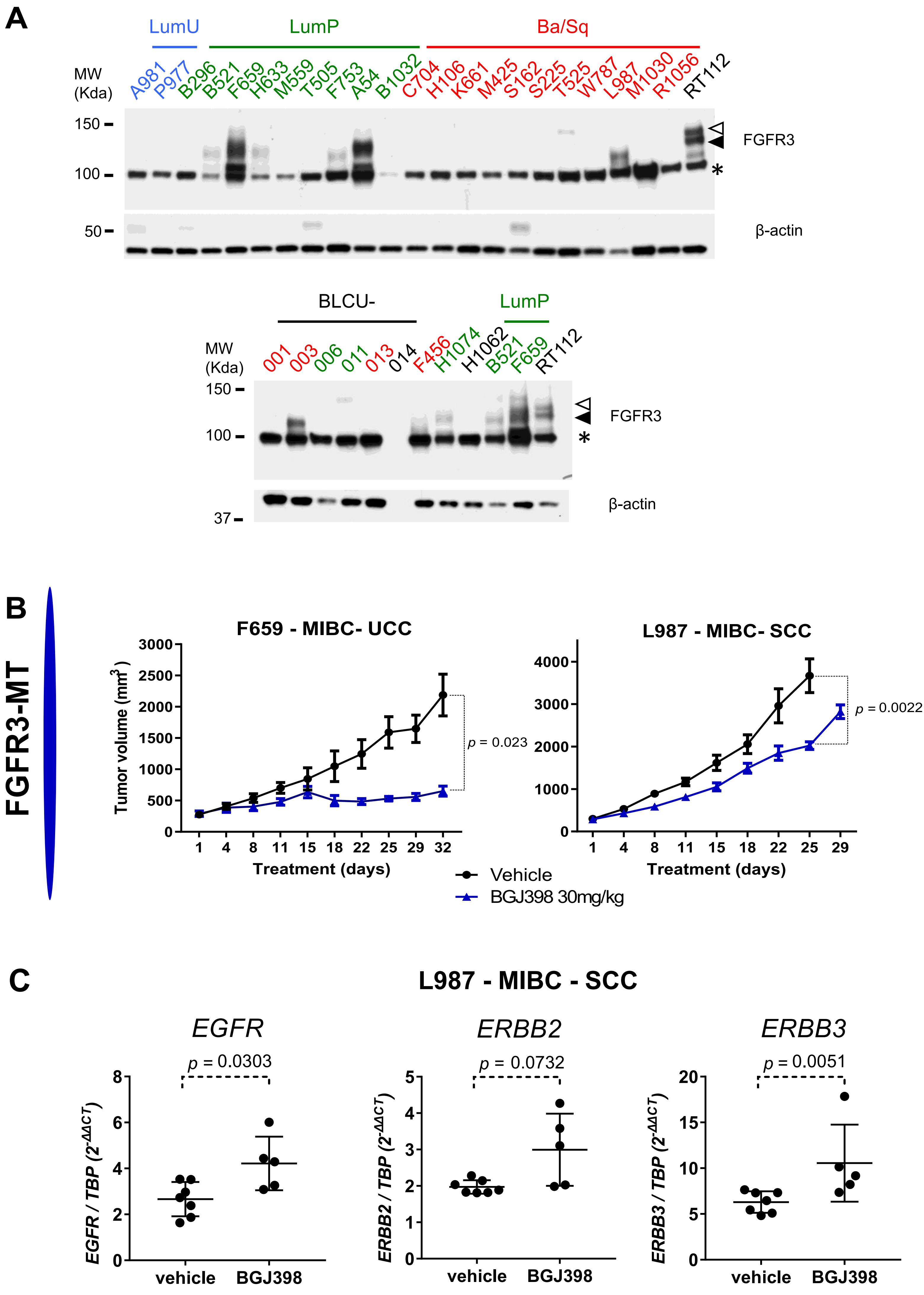
