## Supplemental Method Dragon for "Integrated molecular and pharmacological characterization of patient-derived xenografts from bladder and ureteral cancers identifies new potential therapies"

Technology and analysis pipeline information

**3.1 DRAGON (Detection of Relevant Alterations in Genes involved in Oncogenetics by NGS) analysis pipeline**

In order to assess mutations, copy number alterations, MSI status and tumor mutation burden for routine diagnostic and clinical trials use, a dedicated pan-cancer capture targeting **489 full coding sequence genes**, microsatellites, 3-alleles Single Nucleotide Polymorphisms (SNPs), specific regions of interest in 82 additional genes and a backbone was designed (representing 2.87Mb among which 1.59Mb are coding, see “dragon_v2.bed”).

DNA samples were sequenced using **SureSelect XTHS** Target Enrichment System for Illumina Paired-End Multiplex Sequencing Library on both next generation NextSeq500 & NovaSeq sequencing systems, resulting in the production of 100bp paired-end reads per sample and a 10bp read containing the corresponding molecular-barcoded adaptors (Unique Molecular Identifier per DNA fragment).

Raw sequencing data (BCL) were demultiplexed into 3 reads (Fastq) with the adequate bases-mask information (read and UMI sizes) using *bcl2fastq* (Illumina, v2.20). A first quality control was performed using both *FastQC* (v0.11.5, Andrews, 2010) and *MultiQC* (v1.7, Ewels, 2017) to assess whether all samples were extracted correctly without any quality issue.

These files were then used for two different strategies: i) small, intermediate and largescale variant detection without considering UMIs and ii) verification of a small variant presence after **UMIs deduplication**.

**3.2 Reads mapping**

In the first analysis part, reads were mapped using *BWA mem* (v0.7.15, Li, 2013) on the Human reference genome (hg19 assembly) using default parameters. As a second quality control, statistics regarding the mapping (percentage of aligned reads total and falling into the capture, percentage of PCR duplicates) and the capture coverage were produced using a combination of *SAMtools flagstat* (v1.9, Li *et al*, 2009), *BEDtools coverage* (v2.21.0, Quinlan, 2010) and *PicardTools MarkDuplicates* (v2.6.0, http://broadinstitute.github.io/picard/).

**3.3 Copy Number Alteration**

Copy number alterations were called using the combination of homemade R (v3.2.0, R Core Team, 2015) scripts and facets package (v0.6.0, Shen and Seshan, 2016) with a mimimum mapping quality of 0 and a minimum base quality of 15. Facets performs joint segmentation of total- and allele-specific read counts, and integer copy number calls corrected for tumor purity and ploidy. It only accepts a pair of tumor/normal samples: the normal sample for SHIVA02 patients corresponds to a defined sex-specific unmatched-germline control previously sequenced using the same panel.

- **3.3.1 Genomic profile**

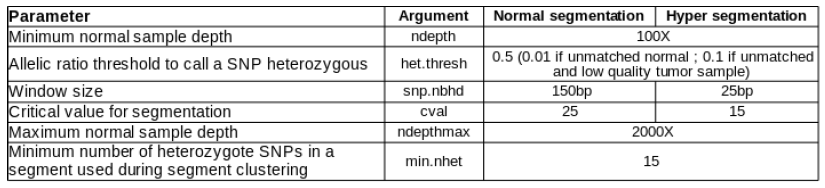
Two rounds of segmentation (a normal step and a hyper-segmented step allowing to recover small focal events) were performed using facets and the following fitted parameters:

Low quality samples (presenting warning with an “het.thres” parameter at 0.01 were fitted again with a more stringent value, 0.1, to avoid hyper segmentation of the BAF profile).

For each segmentation profile, a copy number event was stated per segment depending on the number of copies compared to the estimated ploidy of the sample:

Loss-of-Heterozygosity (LOH) was extrapolated from *facets* genotyping results: segments presenting only one allele (*e.g.*: AA instead of AB for a diploid segment) were considered as LOH.

Focal (<10Mb) amplification and homozygous deletion were extracted from the hypersegmented profile and aggregated with the normal profile. Resulting segments were then annotated with the targeted genes of the panel (see *_cnv_report.tsv).

A whole genome view of the i) B-Allele Frequency (BAF), ii) tumor/normal depth ratio and iii) total (in black) and minor (in grey) copy number profiles can be found in the corresponding graph. In order to help the identification of putative event of interests, **a list of genes was previously created per patient pathology** in order to highlight them successively in orange and green on the first 2 profiles (BAF and depth ratio), interdependently of their copy number status. On the third profile, **only genes presenting focal amplification for a list of oncogenes or focal homozygous deletion for a list of tumor suppressor genes** are highlighted in, respectively, blue and red.

Segments above a log depth ratio of 4 or a total copy number of 20 are limited to these respective values for graphical reasons.

- **3.3.2 Intragenic deletion**

In order to see exon-size intragenic rearrangement, per-sample read depth at extrapositions (every 50bp on a list of priority 1 genes) were added to the list of polymorphisms used as input by facets. After normalizing against control using previous criteria (normal segmentation), the resulting depth ratio per gene (limited to 2 for graphical issues) and per sample were represented as red points using bioconductor R package Gviz (Hahne and Ivanek, 2016) and the distribution of the other samples of the run were displayed as grey boxplots. Noisy samples (presenting an average depth ratio below 1% quantile of the other samples) were discarded from these distribution per gene. A red rectangle helps visualize putative intragenic deletion by highlighting regions of the sample of interest which are below the 2.5% quantile of the other samples.

**3.4 Variant calling**

- **3.4.1 Single Nucleotides variations and small insertions/deletions**

Variant calling of both single nucleotide variations (SNVs) and small insertion/deletions (indels) was then performed on the processed alignment files using a combination of the mpileup module of SAMtools - taking into account anomalous read pairs, without base quality recalibration, considering a minimum mapping quality of 0 and base quality of 15, and a maximum read depth of 1M – and VarScan2 mpileup2cns (v2.4.3, Koboldt et al., 2009). Variants were reported if the number of reads supporting the alternative allele was superior or equal to 2, at a locus covered by at least 10 reads, and if the allelic ratio of this variant was superior or equal to 1%. When available, Varscan2 somatic mode was used to call variants using both tumor and matched-germline samples with similar parameters. A bug of VarScan2 mpileup2cns prevents SNVs presenting multi-allelic positions to be called: mpileup2snp was therefore used to catch these alterations that were then added to the variants call using an in-house R script.

*Bcftools* (v1.3.1, Li *et al.,* 2011) “norm” module was then used to split multi-allelic variants into several rows and to left-align and normalize indels. Using a suite of in-house *python* (v2.7.13) scripts, variants were finally annotated with different metrics regarding variants context inside the reads (depth of coverage, strand bias, median base qualities, variant position...).

- **3.4.2 Intermediate-size indels**

A dedicated strategy using *transIndel* (v0.1, commit 7098bd6, Yang, 2018) was applied to call intermediate-size indels, with the following parameters: minimum mapping quality of 0, read depth above 10X, length between 10bp and 1000bp, covered by at least 30X and presenting an allelic ratio above 1%.

- **3.4.3 Annotation**

Annotations from several databases (RefSeq, dbsnp v150, COSMIC v86, 1000g project 08/2015 version, ESP6500, gnomAD (all and ethnies), ICGC v21, and dbnsfp v35 predictions) were provided by **Annovar** (04/16/2018 version, Wang et al.¸ 2010) to annotate small variants. Only the RefSeq database was used for intermediate-indel. During this step, all variants present in -10/+10bp of each exon junction were defined as splicing.

Finally, a suite of in-house *python* scripts were used to add supplementary annotations (correct HGVS nomenclature (*python* hgvs package v1.2.5 and 2018 UTA database), MaxEntScan, Grantham and ESR scores) and to format the variant table as requested (see “*_table_report.tsv”).

- **3.4.4 UMI**

All variants (SNVs and small indels) called per sample were looked at in the alignment files containing consensus UMIs using an in-house python script (check_variants) with the same filtering strategy: mapping quality of 0 and base quality of 15. Allelic ratio of the variants and depth of overage of the loci after UMI deduplication were then added to the table report previously generated. At this time, the **use of UMI in the analysis is only descriptive.**

- **3.4.5 Variant prioritization**

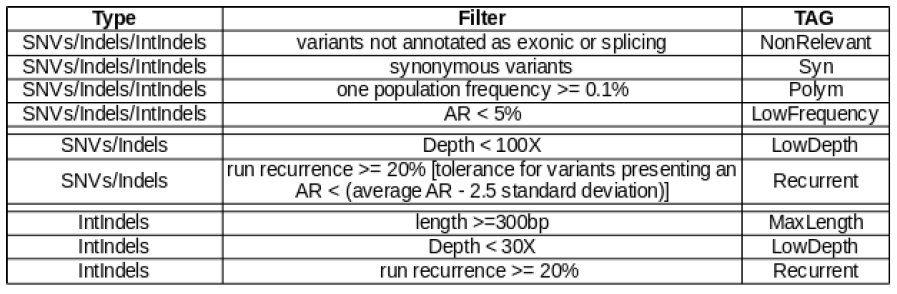
Variants were tagged depending on the following criteria:

Variants **passing these filters were tagged as “PASS”**. A list of defined noncoding regions of interest and coding hotspots were always considered “PASS” independently of the filtering strategy.

When the matched-germline was available, a somatic variant calling was performed allowing to know the allelic ratio of the variant in the germline sample. An additional tag “TAG_MATCH” categorizing variants as “Somatic” if the germline allelic ratio was below 0.5%.

Finally, an additional “TAG_UMI” was added to describe “Low_Freq_Umi” (allelic ratio with UMI < 5%) and “Low_Depth_UMI” (depth with UMI <30X).

All variants (SNVs, indels, intermediate indels) not tagged as “*NonRelevant*”, “*Syn*” or

“*Polym*” were reported in a dedicated table (see “*.variants_report.table1.tsv”).

- **3.4.6 Tumor Mutational Burden (TMB)**

Tumor mutational burden is a measurement of mutations per megabase carried by tumor cells and is a predictive biomarker being studied to evaluate its association with response to immunotherapy. The difficulty consists of identifying the somatic alterations in the absence of the matched-germline sample. Furthermore, the type of variants to consider remains unclear.

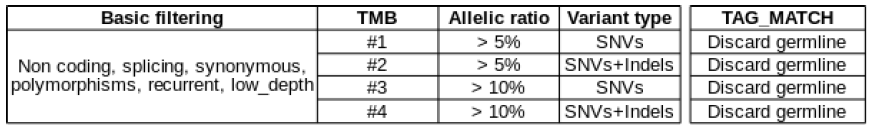
Several TMB values were computed per sample. Coding variants (at the exceptions of splicing ones, therefore exons-only which represent 1.59Mb), without synonymous nor polymorphisms (0.1% MAF) or recurrent variants, and covered enough (not tagged as Low_Depth) were considered in all those calculations. 3 factors may vary: allelic ratio, variant type and if there is a matched-germline (see “*_tmb_results.tsv”). Previously “Saved” variants were not considered in the TMB.

**3.5 Identitovigilance**

In order to identify putative samples swaps or mismatches, a hierarchical clustering was generated using the variants state in all samples (mutated or not mutated) using the Jaccard distance and the Ward linkage function (vegan v.2.5-5 - R package v.3.6.0). Samples are annotated by all “a priori” informations filled in the SampleSheet and an additional quality annotation is added to identify samples that might cluster together due to low quality (average depth of coverage below 100X).

**3.6 MicroSatellites Instability (MSI)**

Given tumor only sequencing data, *MSIsensor2* (commit ebdbf42, niu-lab), uses machine learning models to figure out the MSI status for a distribution per microsatellite. Considering all loci (microsatellites larger than 3bp, homopolymer size between 5bp and 50bp) covered by at least 1X, **percentages of unstable loci (score)** were computed per sample (see “*_msi.tsv”).

Given a pool of MSI-high and MSS samples as controls, unstable loci were represented on an oncoPrint using *R* package *ComplexHeatmap* (v2.0.0, Gu *et al.,* 2016).

**3.8 Coverage quality control**

A more detailed notion of the percentage of each gene, per barcode, **covered by at least 100X**, in the processed alignments, was also provided using a combination of *awk*, *SAMtools mpileup, BEDtools intersect, multiinter and merge* (see “*.coverage_percentage.tsv”). Bases covered by less than 100X were reported per barcode using the same strategy (see “*.coverage_maker.tsv”). Genes belonging to the patient pathology were tagged in these two

files to facilitate the search for genes of interest that might be badly covered.

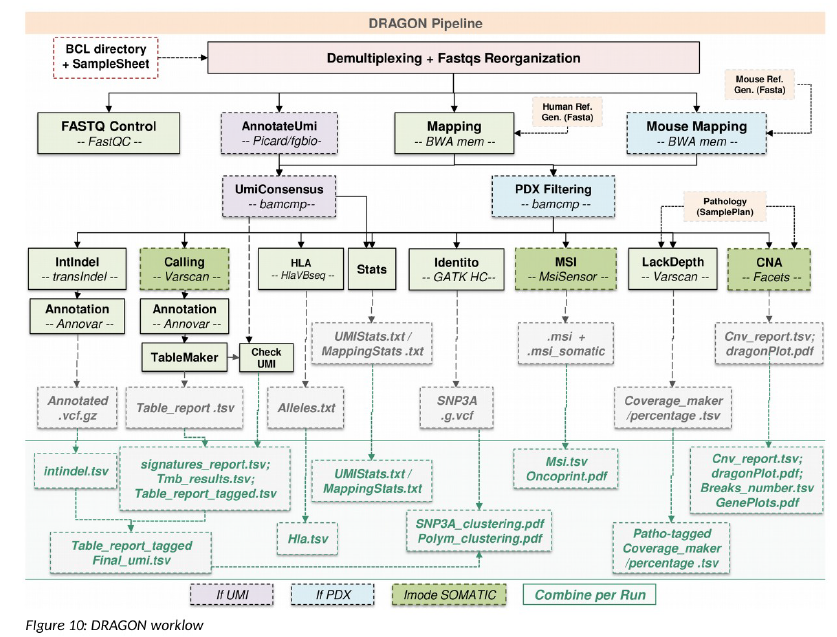

***Figure 1: DRAGON workflow***

**4. Decision strategy in SHIVA02 clinical trial**

**4.1 Dragon**

The Next-Generation sequencing (NGS) strategy used here is dedicated to prepare indexed paired-end libraries of tumor DNA using the Agilent Sureselect XT-HS library prep kit. The kit supports sequencing targeted regions of the genome spanning 2.7 Mb and 571 genes distributed along 23 chromosomes. This panel is called DRAGON (Detection of relevant alteration in genes involved in oncogenetics by NGS).

Briefly, between 50 ng of input DNA are used to build the libraries according to manufacturer’s protocol. The pool is finally sequenced on a NovaSeq Sp 2x100 bp flowcell.

**4.2 Variant classification**

Variants will be classified into 5 classes:

Class1	Pathogenic variant (known hotspot variants)

Class2	Likely pathogenic variant (based on in silico analysis)

Class3	Variant of uncertain significance

Class4	Likely benign variant (probably no pathogenic based on in silico analysis)

Class5	Benign variant (other variants certainly no pathogenic)

**4.3 Activating alterations in oncogenes**

- **4.3.1 CNV (copy number variation)**

***Focal amplification***

**- fold change ≥ X8 and/or LogRatio ≥ 2.00**

- if the LogRatio stands between 1.5 and 2, the copy number status of the locus will be assessed by the biologists by taking into consideration the percentage of tumor cells in the sample analyzed, the ploidy, and the minimum size of the amplification when available.

- confirmation by FISH will not be performed in all cases

***Intragenic rearrangement***

- NGS strategy used is not able to accurately detect the intragenic rearrangements

- **4.3.2 SNV (variant)**

***Missense hot-spot mutations***

Will be taken into consideration based on the literature, the prediction database (PolyPhen-2 (build r394 of v2.2.2 and consecutive versions), SIFT (SIFT 6.2.1 and consecutive versions), etc …), localization of the variation (i.e. does the variant stand in a functional domain, public tumor database (COSMIC (v81 released 09-MAY-17 and consecutive versions), cbioportal (Version 1.7.1 and consecutive versions), etc …) and local variant database.

***In frame Ins/Del***

Will be taken into consideration if described in public tumor database (Cosmic, tumorportal, etc …) or in the literature (ex: deletion in ex19 of EGFR).

***Splice site mutation***

Will be taken into consideration if the resulting transcript stays in frame and if described in the literature (ex: mutation involving the splicing of ex14 of MET).

***Allelic frequency***

molecular biomarkers predictive to targeted therapy sensitivity ≥ 5%

molecular biomarkers predictive to targeted therapy resistance ≥ 1%

***Variant not considered as activating***

- Polymorphism with a frequence > 0.1% in 1000genome (08/2015 version) and/or ESV (ESP6500)

- Synonymous (but not in the first/last 10 bp of the exons since they can alter the splicing)

- Nonsense mutation

- Frameshift

**4.4 Inactivating alterations in tumor suppressor genes**

Only bi-allelic alteration will be considered:

•	Homozygous inactivating alteration

•	Composite heterozygous inactivating alteration

•	Inactivating mutation + LOH

•	Homozygous deletion

***Variants considered as inactivating***

- Nonsense mutation (except rare cases as those located at the end of the gene)

- Frameshift Ins/Del

- Splicing mutation that result in a frameshift

- Intragenic large deletion or duplication

- Class1 missense mutation if already described in the literature as inactivating

***Variants not considered as inactivating***

- Polymorphism with a frequence > 0.1% in 1000genome and/or ESV

- Silent (but not in the first/last 10 bp of the exons since they can alter the splicing)

- Unknown missense mutation

- Frameshift
